# Genetic structure and genome-wide association study of the traditional Kazakh horses

**DOI:** 10.1101/2023.03.29.534422

**Authors:** A. Pozharskiy, A. Abdrakhmanova, I. Beishova, A. Shamshidin, A. Nametov, T. Ulyanova, G. Bekova, N. Kikebayev, A. Kovalchuk, V. Ulyanov, A. Turabayev, M. Khusnitdinova, K. Zhambakin, Z. Sapakhova, M. Shamekova, D. Gritsenko

## Abstract

Horses are traditionally used in Kazakhstan as a source of food and as working and saddle animals as well. Here, for the first time, microarray-based medium-density SNP genotyping of six traditionally defined types and breeds of indigenous Kazakh horses was conducted to reveal their genetic structure and find markers associated with animal size and weight. The results showed that the pre-defined separation between breeds and sampled populations was not supported by the molecular data. The lack of genetic variation between breeds and populations was revealed by the principal component analysis (PCA), ADMIXTURE, and distance based analyses, as well as the general population parameters expected and observed heterozygosity (*H*_*e*_ and *H*_*o*_) and between group fixation index (*F*_st_). The comparison with previously published data on global horse breed diversity revealed the relatively high level of individual diversity of Kazakh horses in comparison with the wellknown foreign breeds. The Mongolian and Tuva breeds were identified as the closest horse landraces, demonstrating similar patterns of internal variability. The genome-wide association analysis was performed for animal size and weight as the traits directly related with meat productivity of horses. The analysis identified a set of 60 SNPs linked with horse genes involved in the regulation of processes of development of connective tissues and the bone system, neural system, immune system regulation, and other processes. The present study is novel and introduces Kazakh horses as a promising genetic source for horse breeding and selection.

## Introduction

Horses (*Equus caballus* L.) have traditionally played a central role in the whole history of Kazakhstan and the Kazakh people. The eneolithic Botai culture (Northern Kazakhstan) contains arguably the earliest evidences of the use of horses by the local tribes (Levine, 1999), however, it remains disputable whether horses were domesticated or obtained by hunting (Outram et al., 2009, 2021; Taylor and Barrón-Ortiz, 2021). Genomic data revealed that Botai horses were closer to Przewalski’s horses than to modern domestic lineages (Gaunitz et al., 2018), thus, even if the Neoilthic horse domestication had taken place at Botai, it had occurred independent of the main course of horse domestication (Kyselý and Peške, 2022). Nevertheless, horses became an important part of the steppe pastoralism and nomadism at the area of modern Kazakhstan and Central Asia as early as the Bronze Age (Frachetti and Benecke, 2009; Outram et al., 2012). From the SakaSkythian tribes to three Kazakh Hordes, throughout the process of the Kazakh ethnogenesis, horses were not only an essential economical resource for the local Nomadic peoples (Chang, 2015), but also became an important part of the cultural legacy inherited by independent Kazakhstan, with rich symbolic connotations and influence on Kazakh language (Sarbassova, 2015).

Traditional Kazakh husbandry practices have not changed significantly for centuries and have been based on grazing and seasonal transhumance (migrations between *zhaylau* and *qystau*, summer and winter pastures, respectively). The conditions of horse pasturing have been kept close to natural: free grazing under the herdsman’s control; in winter, horses find their fodder from under the snow—*tebindeu*. Herds of horses were the main measure of one’s wellness; the owners selected the best animals or interchanged them with other breeders to keep and improve the horses’ valuable qualities, such as strength, endurance, and the most important for nomads, meat and milk productivity. As a result of such folk selection over hundreds and thousands of years, the Kazakh horse was formed, with some traditional types distinguished based on their qualities and geographical distribution, the most known of which are Zhabe^1^, Adai^2^, and Naiman horses. Zhabe and Adai horses are the most common types. The Zhabe type is specialized for meat and milk production, and the Adai type has more expressed qualities of saddle horses; both these types originated in Western Kazakhstan and are used in all regions of the country (Dmitriev et al., 1989). The third type, Naiman, has been traditionally bred by the inhabitants of Dzhungar Alatau as the universal horses for use in mountainous conditions; this type is similar to Mongolian horses and considered the most subtle of Kazakh horses (Dyussegaliyev, 2022). All these types are well adapted for the traditional Kazakh methods of seasonal pasturing and transhumance (Barmintsev, 1958).

Traditional Kazakh horses became a progenitors of new breeds by crossing with the animals with desirable traits. Kostanay^3^ breed was under development since 1887 year by crossing Kazakh mares with stallions of saddle breeds, Don, Astrakhan, Strelets, Thoroughbred, to combine their best qualities (Dmitriev et al., 1989). The breed was finally registered in 1951 (Nechayev et al., 2005), and work on its improvement has continued into the present. These horses have good saddle and draft characteristics and are suitable for both stable and steppe maintenance (Dmitriev et al., 1989).

The Kushum horses were bred between 1931 and 1976 by crossing Kazakh horses with Thoroughbreds, Don breeds, and their half-breeds, initially as military horses and later, after World War II, as steppe herd horses for meat and milk production. The resulting breed is versatile; the horses have high endurance and are capable of high live weight gai (Dmitriev et al., 1989).

The Mugalzhar breed was developed based on Kazakh horses between 1969 and 1998 to improve meat and milk productivity (Iskhan et al., 2019). Non-specific crossings and selection within the Zhabe breed allowed a significant increase of live weight (80–100 kg for mares, 100–120 kg for stallions, compared to the original horses), without changing technology of their maintenance (Dyussegaliyev, 2022).

A significant damage to the Kazakh horse population was caused by the Soviet policy of the involuntary collectivization and eradication of private animal ownership, and during the period between 1928 and 1958, the amount of Kazakh horses was reduced from 4,640,000 to 300,000 (Nechayev et al., 2005). According to the data of the Bureau of National statistics of the Agency for Strategic planning and reforms of the Republic of Kazakhstan (Bureau of National statistics, 2023), the total amount of horses (including foreign breeds) in the country was 1,666.4 thousands in 1991; after the drastic decrease in the 1990s, the number of horses was growing steadily and reached 3,489.8 thousands in 2021, due to the development of husbandry in Kazakhstan.

Kazakhstan is the second large horse meat producer in the world after China, however, it is mainly limited to the domestic market, as the country is not a significant exporter of horse meat (Jastrzębska et al., 2019). With the growing interest in horse meat as a safe and nutritious alternative to beef, despite a prejudicial attitude existing in many countries (Stanciu, 2015), Kazakhstan has a potential to become an important provider in the global horse meat market. This requires an extensive modernization of horse breeding to comply with the internationally acclaimed standards. An important aspect of such modernization is a wide implementation of the contemporary methods of molecular genetics and genomics in breeding practices to better understand the genetic structures of horse lines and breeds, improve classification and management of horse genotypes, assist selection using molecular markers associated with valuable traits, etc. The commercial microarray genotyping panels for animals contain tens or hundreds of thousands of SNP markers selected to reflect the total genetic variability, helping scan genomes for potentially important polymorphisms without expensive whole-genome sequencing. In Kazakhstan, SNP microarray genotyping was previously used to describe genetic structures of the local breeds of sheep, another animal of essential importance for the country (Pozharskiy et al., 2020; Zhumadillayev et al., 2022). For horses, the EquineSNP50 panel was developed and proven to be suitable for genome-wide association analysis and studies of horse diversity (McCue et al., 2012; Petersen et al., 2013a; b).

The scope of the present work was to explore genetic diversity of Kazakh horses using the Equine80k SNP microarray, the upgraded and expanded version of the EquineSNP50 array. The horses of three traditional types (Zhabe, Adai, and Naiman) and three derivative breeds (Kushum, Kostanay, and Mugalzhar) were sampled from herds in different regions of Kazakhstan to reveal their genetic variability between and within breeds. The previously published data of genetic diversity of horse breeds from all over the world was also used for comparison. Also, a genome-wide association study (GWAS) was performed to find SNPs associated with size and live weight of the horses. This is the first study involving high-density SNP array into an investigation of traditional Kazakh horses in Kazakhstan. The obtained results will increase understanding of the genetics of Kazakh horses and their place in the global diversity of horse breeds and lay a basis for further molecular and genomic studies.

## Materials and methods

### Sample collection and data acquisition

Genetic materials of three traditional types and three breeds of Kazakh horses (for convenience, we will further refer these as six breeds) were collected at 25 horse farms in different regions of Kazakhstan (Figure 1; Supplementary table 1). Hairs were sampled from horses’ tails and/or manes and stored at +4°C until further use; the hair follicles were used for DNA extraction. DNA was isolated using the kit “DNK-Extran2” (Syntol, Russian Federation) following the manufacturer’s protocol and quantified using Qubit 4 Fluoremeter with the Qubit dsDNA Broad Range reagent (Thermo Fisher Scientific, USA) for downstream SNP genotyping.

**Figure 1.**
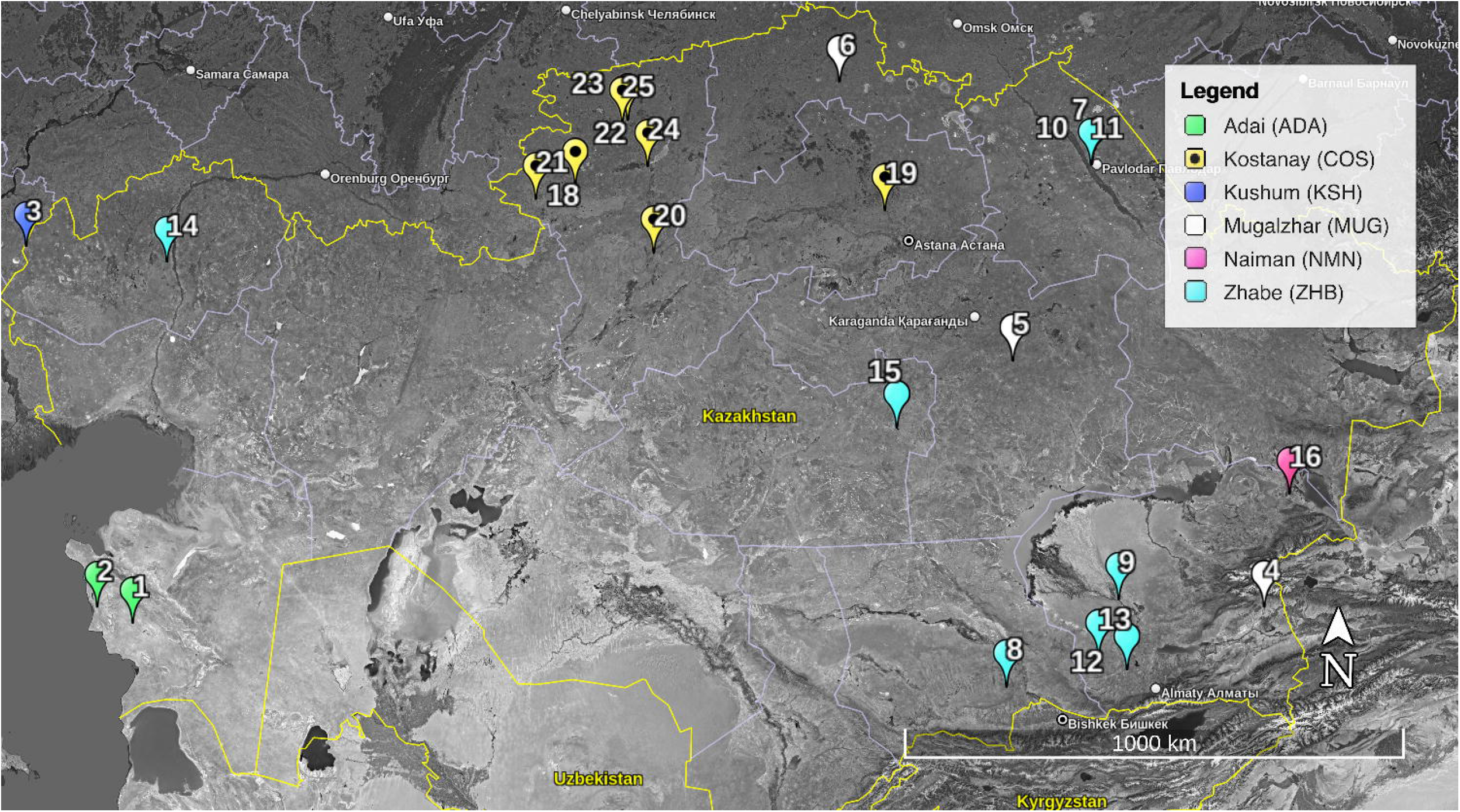
Geographical distribution of horse farms sampled in the study. 1 Kuanysh, 2 Kozhyr-Ata, 3 – Alem, 4 – Orazbay, 5 – Bakytbek, 6 – Kara-Tomar, 7 – Agro-Damu, 8 – Kalka, 9 – Zhaksylyk, 10 – Anar, 11 – Akzhar, 12 – Sayahat, 13 **–** Kyzylsok, 14 – Ernur, 15 – Orazaly, 16 – Tok-Zhailau, 17 – Kazakh Tulpary, 18 – ChV, 19 – Tasmukhambetov, 20 – Kostanay Tulpary, 21 – Stepnoye, 22 – Kabazhaev, 23 – Baranovskiy, 24 – Akhmetov, 25 – Zhussupov.

The measurements of horses were taken prior to hair sampling. The measurements included the height at the withers, oblique body length, chest circumference, cannon bone circumference, and body weight.

SNP genotyping data on the genetic diversity of foreign horse breeds (Petersen et al., 2013a) was retrieved from the Open Science Foundation repository (https://osf.io/gx42p/, accessed 20.07.2022), with the explicit permission from Dr. Jessica Petersen.

### SNP genotyping and quality control

Genotyping was performed using Equine80k SNP array with the iScan system (Illumina, USA) according to the manufacturer’s protocol. The assignments of genotypes and primary quality control were conducted using the Genotyping module of the GenomeStudio software (Illumina). The data was filtered using the following thresholds (the primary quality control): call rate ≥ 0.9, median GC score ≥ 0.8 for samples; call frequency ≥ 0.95 and GT score ≥ 0.7 for SNP. The indel markers presented in the array were also excluded. All data satisfying these criteria were exported and transformed to PLINK text input files (.ped + .map) using general data handling utilities of R (R Core Team, 2019). PLINK1.9 (Purcell et al., 2007; Chang et al., 2015) was further used to exclude SNPs with minor allele frequencies ≤ 0.05 and those deviating from Hardy-Weinberg equilibrium with a *p-*value threshold of 1·10^−10^. The missing genotypes were imputed using BEAGLE (Browning et al., 2018).

### Genetic structure analysis

All samples were tested for the internal and between-population structure using ADMIXTURE (Alexander and Lange, 2011) for K from 1 to 10 with 10 cross-validation replicates. Based on the results the outlier genotypes were identified and taken into account for any further analyses.

The general population genetic analysis was performed using PLINK1.9 and summarized using general R functions. It included the evaluation of linkage disequilibrium (LD), expected and observed heterozygosity (*H*_*e*,_ *H*_*o*_), and between-population fixation index (*F*_*st*_). The analysis was performed for the whole sample for breeds and populations. Only populations with more than 50 sampled individuals were considered for population-based analysis. The markers in the strong LD (*r*^*2*^ > 0.7) were not considered for the population statistics.

The comparative analysis of the Kazakh horse genotypes with respect to the well-known foreign horse breeds was performed using data from (Petersen et al., 2013a). The dataset was transformed to fit the data format used throughout the present work using general R scripting tools and merged with our data. The merged dataset included 10 randomly selected individuals from each local population with more than 10 samples and all individuals from populations of the lower sample size. Additionally, the outlier genotypes were added to identify the possible sources of admixture. The merged dataset was finally filtered to include only SNPs with call frequency ≥ 0.95. A principal component analysis (PCA) was performed using PLINK and visualized using R with the ‘ggplot2’ package (Wickham, 2016). The ADMIXTURE analysis was run for K from 1 to 40, with 10 crossvalidation replicates and visualized using CLUMPAK (Kopelman et al., 2015). The distance matrix was calculated using Manhattan distance in the ‘dartR’ package, and the neighbor joining tree was constructed using ‘ape’ package and visualized using FigTree (Rambaut, 2018) software.

### Genome wide association analysis

The genome-wide association analysis was performed using PLINK1.9 supplemented with R scripting, including use of the ‘ggplot2’ package to create Manhattan plots.

The results of the genetic structure analysis were taken into account for the sample selection for GWAS. The association analysis was conducted for weight and size. The animals of the age below three years and the outliers were excluded from the analysis. Pearson’s correlation test was used to ensure the independence of the phenotypic variables from age. The size variable was defined using measurements of the height at the withers (HW), oblique body length (OBL), chest circumference (CC), and cannon bone circumference (CBC). These parameters were normalized by subtracting the mean and dividing by the standard deviation, then PCA was performed, and the first component was selected as the new size variable.

The genome-wide association analysis was performed using a linear regression test with the adaptive Monte-Carlo permutation method of *p-*value correction for multiple comparisons (PLINK’s ‘--linear perm’ command). SNPs in strong linkage disequilibrium were excluded from the analysis (*r*^*2*^ > 0.7).

The markers with the resulting corrected *p-*value below the threshold of 0.001 were annotated using the Variant Effect Predictor tool (VEP) (McLaren et al., 2016) and the DAVID web server (Huang et al., 2009; Sherman et al., 2022). The horse genome assembly EquCab3.0 (GCA_002863925.1) was used as a reference for annotation.

## Results

### Genotyping and genetic structure analysis

Samples of six Kazakh horse breeds were collected from 25 livestock farms in the North-Kazakhstan, West-Kazakhstan, East-Kazakhstan, Mangystau, Akmola, Kostanay, Zhambyl, Almaty, and Aktobe regions (Figure 1, Supplementary table 1). A total of 2,200 horses were sampled and processed for DNA isolation and SNP genotyping. As a result of selection based on quality control of extracted DNA samples and subsequently obtained SNP genotypes, 2,020 individuals were retained for the analysis. For 1,876 and 1,883 samples, respectively, measurements of size and weight were available.

A total of 74,116 SNPs for 2,020 samples satisfying the selected primary quality control criteria were obtained. Further filtering based on MAF and HWE left a total of 60,987 markers for further analyses.

All breeds and populations with 50 or more sampled individuals were analyzed using general population statistics, expected and observed heterozygosity, and pairwise F_st_ between breeds and populations (Table 1). All breeds and populations had very close values of heterozygosity; on average, 0.3462 and 0.3432, standard deviations 0.0051 and 0.0043, for expected and observed heterozygosity, respectively. All pairs of breeds and populations demonstrated F_st_ not exceeding 0.001, indicating a very low degree of differentiation between sampled groups. The lowest between-breed value was observed for Kostanay and Adai (F_st_ = 0.0003) and the highest value was between Kushsum and Adai (F_st_ = 0.005).

**Table 1.**
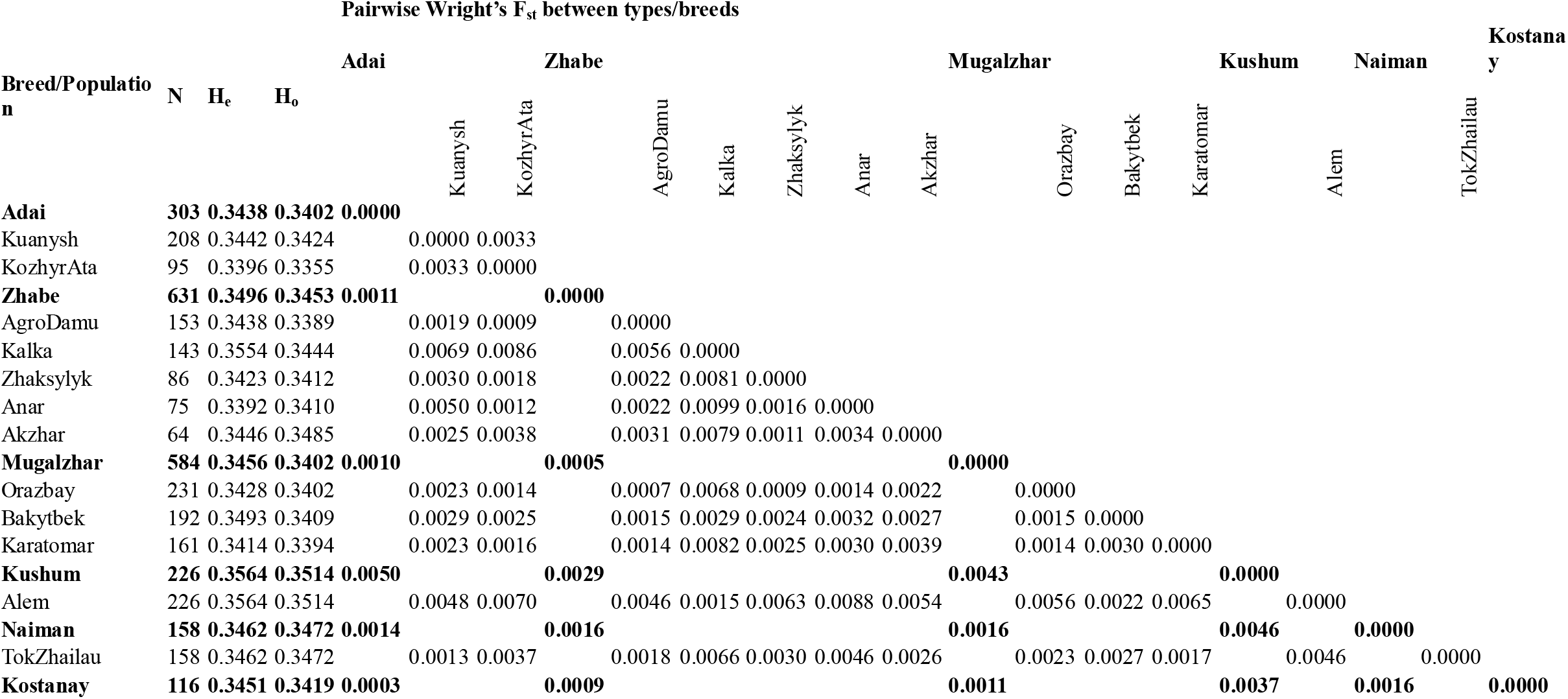
Population genetic statistics of horse breeds and populations. Population with less than 50 sampled individuals are not shown, but accounted for breeds. N – number of samples, H_e_ – expected heterozygosity, H_o_ – observed heterozygosity.

**Table 2.** Foreign breeds used for comparative analysis (data by (Petersen et al., 2013a))

| Abbreviation | Breed name | Country of origin* | Number of samples |
| --- | --- | --- | --- |
| AKTK | Akhal Teke | Turkmenistan | 19 |
| AND | Andalusian | Spain | 18 |
| ARR | Arabian | Arabia | 24 |
| BEL | Belgian | Belgium | 30 |
| CSP | Caspian | Iran | 18 |
| CLYD | Clydesdale | UK | 24 |
| EXMR | Exmoor | UK | 24 |
| FELL | Fell Pony | UK | 21 |
| FIN | Finnhorse | Finland | 27 |
| FLCR | Florida Cracker | USA | 7 |
| FM | Franches-Montagnes | Switzerland | 19 |
| FT | French Trotter | France | 17 |
| HAN | Hanoverian | Germany | 15 |
| ICE | Icelandic | Iceland | 25 |
| LUST | Lusitano | Portugal | 24 |
| MNGP | Mangalarga Paulista | Brazil | 15 |
| MARM | Maremmano | Italy | 24 |
| MINI | Miniature | Unknown | 21 |
| MON | Mongolian | Mongolia | 19 |
| MOR | Morgan | USA | 40 |
| NFST | New Forest Pony | UK | 15 |
| NSWE | North Swedish Horse | Sweden | 19 |
| NORF | Norwegian Fjord | Norway | 21 |
| PT | Paint | USA | 25 |
| PERC | Percheron | France | 23 |
| PERU | Peruvian Paso | Peru | 20 |
| PRPF | Puerto Rican Paso Fino | Puerto Rico | 21 |
| QH | Quarter Horse | USA | 40 |
| SB | Saddlebred | USA | 25 |
| SHET | Shetland | UK | 27 |
| SHR | Shire | UK | 23 |
| STBDNor | Standardbred (Norway) | USA | 25 |
| STBDUS | Standardbred (US) | USA | 15 |
| SZWB | Swiss Warmblood | Switzerland | 14 |
| TB_UK | Thoroughbred (UK/Ire) | UK | 19 |
| TB_US | Thoroughbred (US) | UK | 17 |
| TUVA | Tuva | Russia | 15 |

\* According to the original publication

The analysis of the genetic structure within the whole sample of Kazakh horses with the ADMIXTURE algorithm confirmed the lack of differentiation between breeds and populations. Although the true *K* was not revealed by cross-validation, as the standard cross validation error did not reached the minimum value in *K* runs from 1 to 10, examination of the results allowed us to select *K =* 2 as the optimal structure (Figure 2, a). The results of *K* = 2 demonstrated that the sample was generally homogeneous, with the exception of some outlying genotypes. These outliers were taken into account for further analyses. The further results for *K* from 3 to 10 highlighted within-population variability and did not add information about genetic structure between populations, however the separation of the outliers was supported.

**Figure 2.**
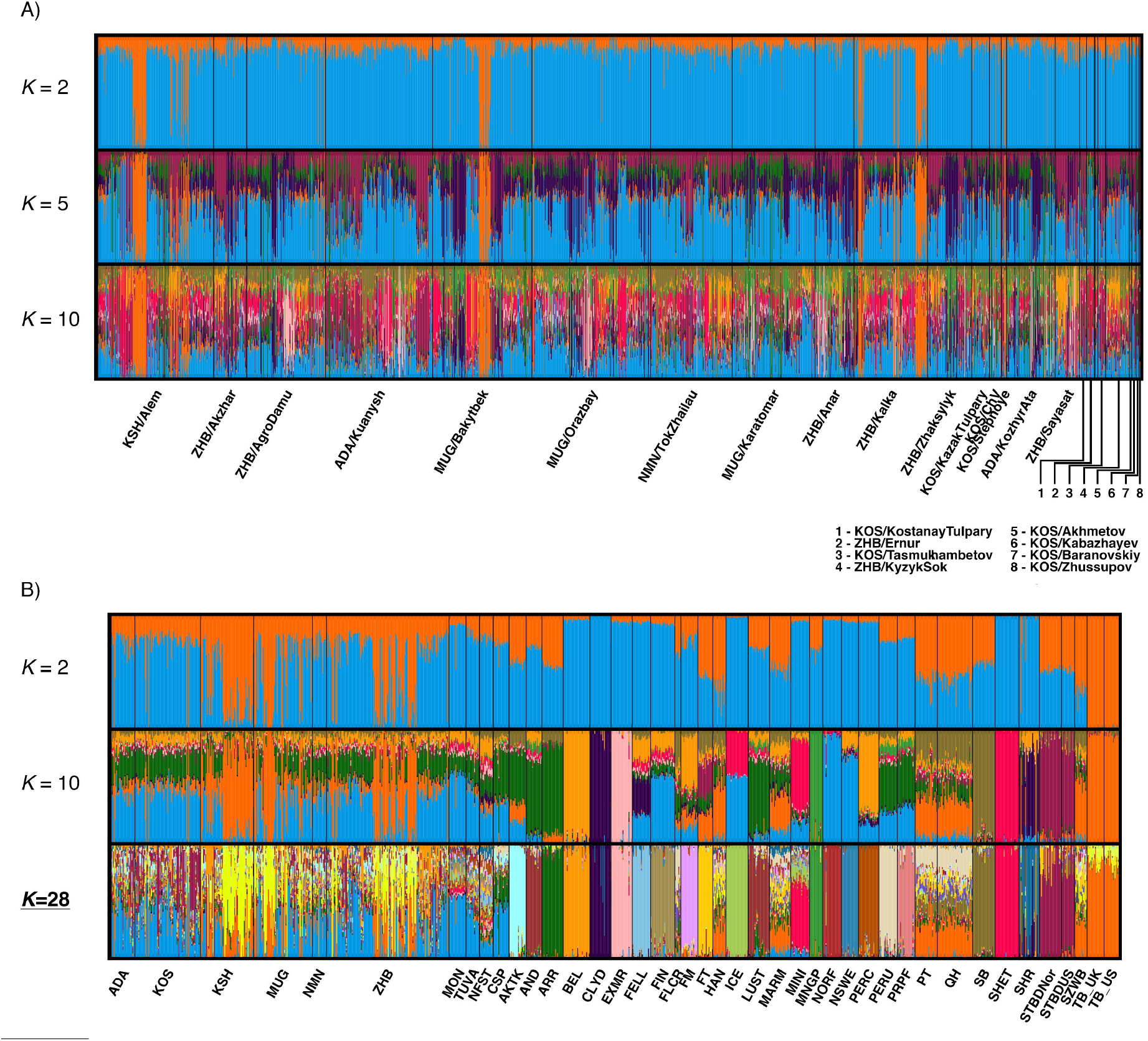
Results of ADMIXTURE analysis. A) complete sample of Kazakh horses of six breeds; B) reduced sample of Kazakh horses in comparison with foreign breeds (data (Petersen et al., 2013a))

Data from Petersen et al. (2013) was used to put the Kazakh horse samples into a context of global horse diversity. As the number of individuals representing the foreign breeds was significantly lower than for Kazakh horse breeds, we achieved balanced data volumes by limiting the number of local horses to not exceed 10 per population; these individuals were sampled randomly. The outlier genotypes mentioned above were additionally selected to identify the probable source of the introgression. The total merged dataset included 35,419 SNPs for 1,176 animals, including 138 Zhabe horses, 74 Kostanay, 66 Mugalzhar, 60 Kushum, and 27 Aday horses. The 10-fold crossvalidation test of the ADMIXTURE analysis identified *K =* 28 as an optimal number of clusters (Figure 2, b). The results demonstrated that Kazakh horses have had higher levels of individual variability compared to well-established foreign breeds. No clustering patterns to distinguished between Kazakh horse breeds were observed. Across the foreign breeds, the most similar to Kazakh horse patterns were observed in Tuva (TUVA) and Mongolian (MON) horses. The identified outlier specimens showed high similarity to the Thoroughbred breed (TB), indicated by orange throughout all *K*’s, however, with the optimal *K* = 28, the outliers of Kazakh horses displayed the high probability of the new cluster (shown pale yellow) with only a minor occurrence in Thoroughbred horses. The results of the PCA (Figure 3) also demonstrated the low level of differentiation between Kazakh breeds. We separately considered the sample with outlier genotypes (Figure 3, b) and without them (Fig. 3, a). Figure 3, a, demonstrates that all Kazakh horses formed one group; the corresponding ellipses and central points were strongly overlapping. In Figure 3, b, the outlier genotypes were shifted towards Thoroughbred horses (populations from the UK and the USA). The breeds closest to Kazakh horses were Mongolian and Tuva horses, as well as the Caspian breed and the groups of Andalusian (AND, Spain) and South American horses. In the overall structure, Kazakh horses had a central position with respect to the directions of distribution of worldwide breeds.

**Figure 3.**
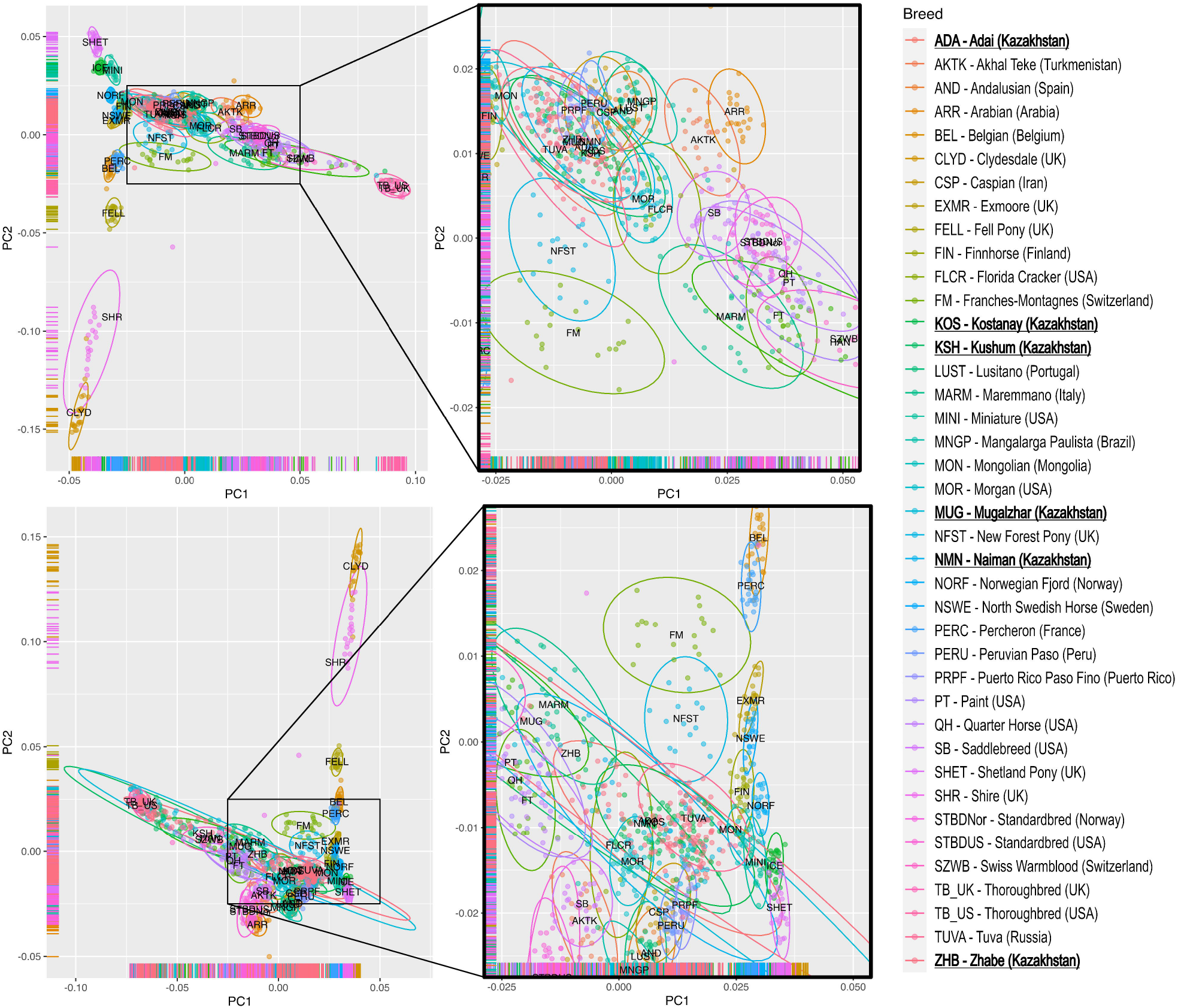
Principal component analysis of Kazakh horses in comparison with foreign breeds. A) outlier genotypes excluded; B) outlier genotypes included. The ellipses indicate the boundaries for points corresponding to breeds, with the confidence level 0.95. The positions of the breed labels correspond to the central points.

In the neighbor joining tree, there were also no clear structures corresponding to Kazakh breeds; most samples were combined into a distinct heterogeneous group, including Mongolian and Tuva horses (Figure 4; Supplementary figure 1). Unlike the results of other methods described above, only a few outlying genotypes diverged from the main group of genotypes. Four horses of the Kostanay, Mugalzhar and Aday breeds were placed close to Thoroughbred horses, and five horses of the Kushum, Mugalzhar, Zhabe, and Naiman breeds were placed between the American breed Morgan and a cluster of South American and Spanish horses.

**Figure 4.**
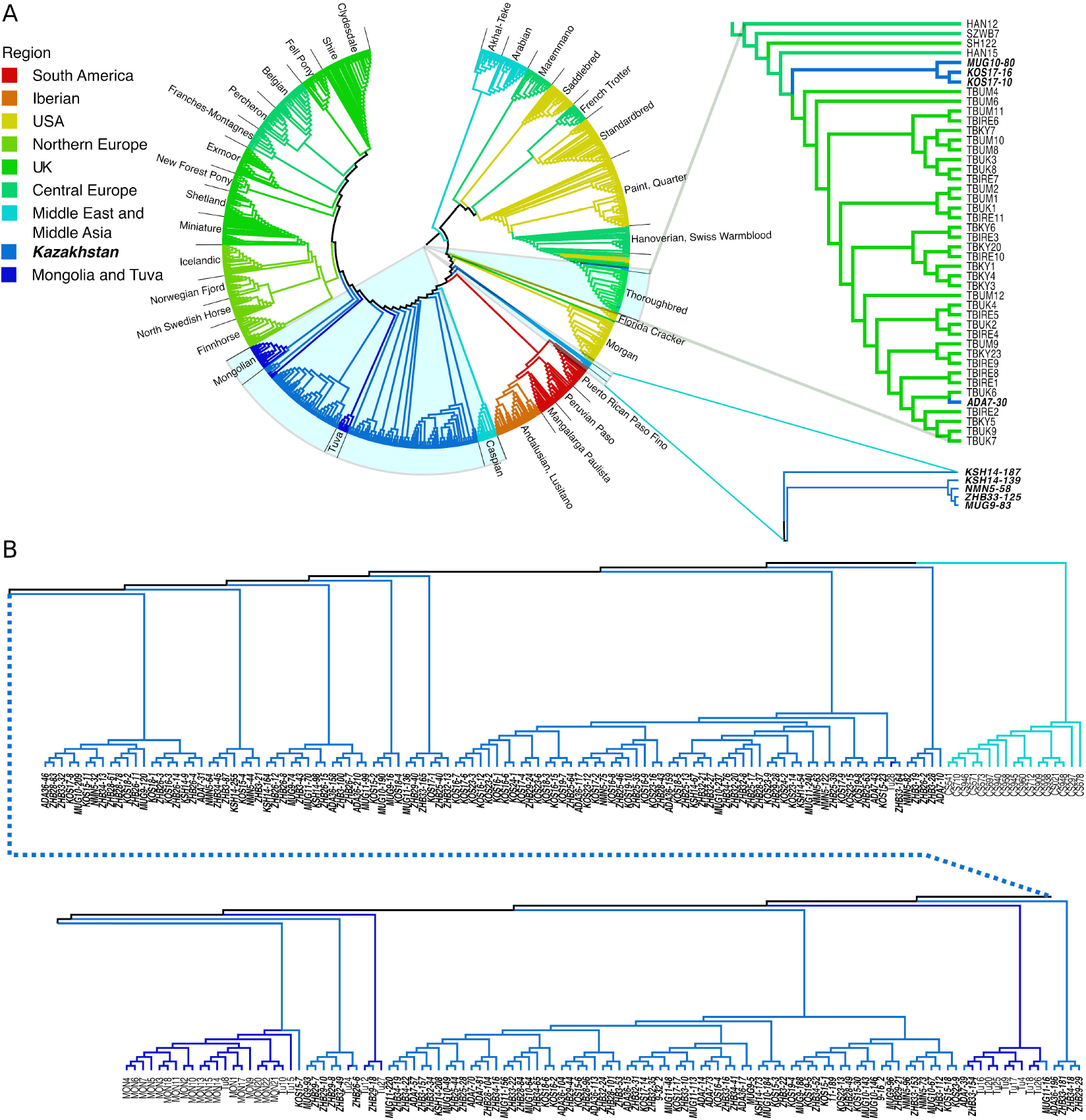
Neighbor joining tree based on Manhattan distances between Kazakh horses and the foreign breeds. A) full view of the tree, with the expanded view of the sub-trees containing outlying Kazakh horse genotypes; B) expanded view of the sub-tree consisting of Kazakh horse breeds.\

### Genome-wide association analysis

The genome-wide analysis of the association between SNP markers with horse body size and weight were performed for all animals with available phenotypic data, excluding horses less than three years of age and the outlying genotypes identified by the genetic structure analysis. The total number of animals was 1,533. To ensure that age did not have an impact as a covariate in the selected sample, all phenotypic variables were tested for Pearson’s correlation against age. The standard body measurements, the height at the withers and oblique body length, displayed no significant correlation at a *p*-value threshold 0.05 (*p*-values of 0.8388 and 0.4211, respectively). Only weak correlations were revealed for chest circumference (0.0841, *p* = 0.000507), cannon bone circumference (0.1011, *p* = 2.922·10^−5^), and weight (0.1121, *p* = 3.496·10^−6^). Four body measurements were combined into a single variable using PCA after normalization; the first principal component describing 81% of total variation was selected as a new size variable.

The association analysis was performed using a linear regression algorithm implemented in PLINK software with the adaptive correction of *p*-values based on a Monte-Carlo permutation test. As a result, a total of 81 and 84 SNPs showed significant associations with size and weight, respectively, at a selected significance level 0.001 (Figure 5). Of these combined SNP sets, 60 variants were associated with the known horse genes using Ensembl’s VEP server and with the corresponding biological processes using DAVID (Table 3, Supplementary table 3). Surprisingly, there was almost no overlap between two sets of markers associated with the respective traits. Only two variants, BIEC2_117960 and BIEC2-187196, showed significant association with both traits. While the former marker was linked with the gene *OR4C269P*, which has had no available gene ontology annotation, the latter was identified in relation to the gene of ecto-5’-nucleotidase (*NT5E*) involved into metabolism of adenosine phosphates. In general, the identified genes play regulatory or signaling roles in a wide range of processes, from the cellular level to the whole organism. The most essential or basic, in our opinion, gene ontology (GO) terms for biological processes (BP) are listed in the Table 4; the complete annotations produced by DAVID are available in Supplementary table 3. The genes *BMP6, DDR3*, and *CREB3L1* are involved in the development and metabolism of connective tissues, including bones. These genes were linked with SNPs significantly associated with weight. The *BMP6* gene had three linked SNPs, which was the highest number across all genes. Several genes, *DPF1, GNAT3, NEGR1*, etc., have been annotated as involved into the development of the neural system. More specifically, the *NEGR1* gene was associated with feeding and locomotory behavior, and the *GNAT3* gene was associated with taste perception. The genes *BMP6, RELA1, AIM2, PDE4D*, and *IGF1R* were associated with regulation of the immune system, in addition to other processes,. The *EIF2AK4* gene was associated with the cellular response to cold stress and protein starvation.

**Table 3.**
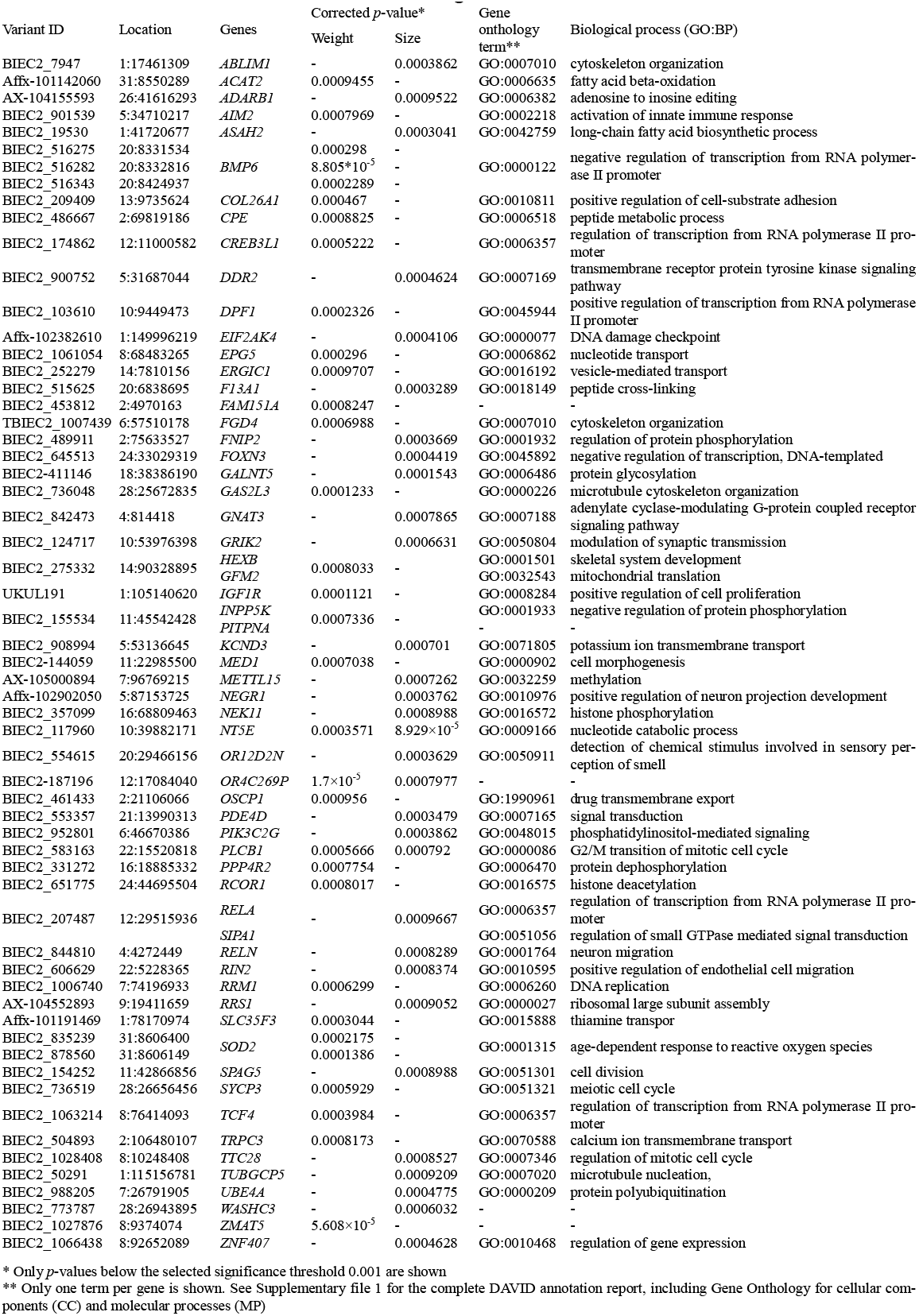
Annotated SNP associated with horse weight and size.

**Figure 5.**
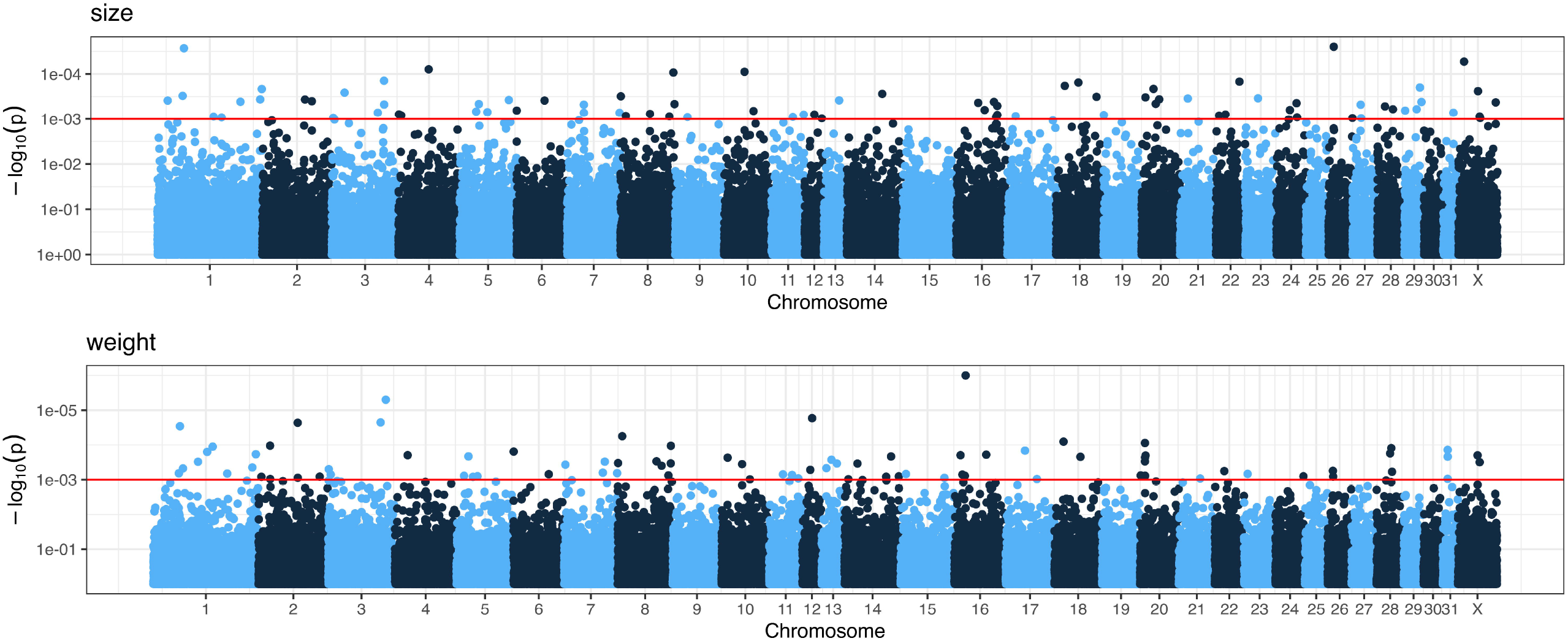
Manhattan plots of *p-*values corrected by the Monte-Carlo permutation test from genome-wide association analysis for horse size (top) and weight (bottom)

## Discussion

This study of Kazakh horses involved three traditional types (Zhabe, Adai, and Naiman) and three relatively recent breeds (Kostanay, Kushum, Mugalzhar), which were derived from the older lines. Except for the Kostanay horses, which are subjected to stable maintenance, control of breeding, and certification of offspring, the other five breeds are being reproduced freely during perennial herd pasturing. However, the genetic data did not support this difference of breeding strategies. All six breeds, including Kostanay, form one group without a notable subdivision corresponding to breeds and populations. This observation was supported by the distance-based and ADMIXTURE clustering methods as well as principal component analysis. Also, pairwise *F*_*st*_ values between breeds and populations were low, indicating almost free gene flow between populations and the factual absence of between-breed boundaries. Hence, we conclude that the studied traditional types and breeds cannot be discriminated from each other on the genetic level, and all Kazakh horses should be considered one ‘breed’. Yet, we question the applicability of the term ‘breed’ to Kazakh horses because of the revealed high level of individual variability. We assume that the Kazakh horses represent a sum of relatively unspecialized genetic lines. In some sense, all Kazakh horses are close to the natural population. Indeed, the traditional Kazakh way of pasturing doesn’t imply strict mating control in herds; gene flow between populations (herds) occurs through trading and exchange of genotypes by horse owners, and the means of modern transporting capabilities reduces the influence of geographical isolation. The close values of expected and observed heterozygosity in breeds and populations of sufficient sizes indicate the absence or low impact of possible selective factors.

Interestingly, the horses of the Tuva and Mongolian breeds (Petersen et al., 2013a)) display similar patterns of individual variability and close proximity to Kazakh horses. According to the published results (Petersen et al., 2013a), these breeds also had higher levels of expected heterozygosity in comparison to many breeds from around the world. Although the small volume of available data for these breeds limits the possible comparisons, we could speculate that Kazakh, Tuva and Mongolian horses could be grouped together. Historically, the nomadic peoples of Central Asia, Mongolia, and Siberia had close interconnections and shared some common ways in horse breeding. Thus, we suggest that the Kazakh, Mongolian, and Tuva horses, and possibly some other breeds not considered here, could be parts of a broader defined landrace, which we would designate as “Nomads’ horses”. In the results of the PCA, all these genotypes had a central position in respect to the main direction of the distribution of breeds. In the study of the Chinese populations of Mongolian horses, a similar PCA pattern was previously observed, with the exception of Kazakh horses (Han et al., 2019). The authors of that study discussed the high homogeneity and the lack of genetic structure between the considered populations and hypothesized that the Mongolian horses may represent the closest descents of the earliest domesticated horse lines. With our results revealing close proximity of Kazakh horses with the Mongolian and Tuva horses, and similar genetic patterns of Kazakh horses, we consider the mentioned study to be in line with our assumption about “Nomads’ horses”. The analysis of horse mitochondrial DNA revealed that, to a significant extent, horses domesticated in the Central Asian region have retained their genetic diversity (Cieslak et al., 2010). We assume that the traditional nomadic ways of horse husbandry played an important role in avoiding a bottleneck effect as they imply only the limited involvement of humans in selection and reproduction.

The analysis of genetic structure of Kazakh horses revealed the presence of significantly deviating genotypes, mainly in populations of Zhabe, Kushum, and Mugalzhar horses. The ADMIXTURE analysis and PCA helped attribute these horses to hybridization with Thoroughbred horses. Indeed, this breed has been popular among breeders in the country, as well as in the world, as the reference saddle and breed horse. Thoroughbred stallions have been imported since the beginning of the twentieth century and are used in crosses to improve the saddle quality of Kazakh horses. However, prior to sampling, all animals used in the work were attested as purebred Kazakh horses of respective breeds and types. This fact raises important questions about the present state of breeding control and certification in Kazakhstan. Our study revealed some significant issues. First, the hybridization of Kazakh horses with other horse breeds, including foreign ones, has been poorly documented, hence the difficulty of tracing back the admixed lineages and their correct identification. Second, the classification of the types and breeds of Kazakh is based on morphological traits and geographical distribution; however, these features do not provide unambiguous boundaries between breeds and, thus, their definition relies mostly on the subjective experience of the breeders.

The lack of genetic structure across the studied breeds and populations allowed us to combine all horses, except for hybrid genotypes discussed above, for association analysis. To date, most genome-wide association studies on horses have focused on racing performance (Littiere et al., 2020; Bailey et al., 2022) and health (Raudsepp et al., 2019). The traits related with food quality of horses have remained out of focus of horse genomics, as horse products, mainly meat, remain exotic or even marginal in many countries (Cieslak et al., 2010). Thus, the study of genetics of such traits is novel, not only for Kazakhstan. Here we tested a set of SNP markers for association to the most general parameters related with meat productivity, live weight and size of an animal. Sixty SNPs were found to be associated with either of these two traits and linked with the functionally annotated horse genes. The set of identified genes included genes involved in various biological processes as regulatory and signal factors. The interesting notion was that almost all significant annotations were related to size or weight independently, despite the obvious correlation between these traits. Among all functionally annotated genes some certain aspects of biological processes potentially related with the traits of interest can be noted. First, the development of connective tissues and bone system, which are crucial for an animal to support its weight and size. Second, the development of the neural system; the more specific effects of the genes *GNAT3* and *NEGR1* on the fodder preferences of horses and so, indirectly, on their growth, could be an interesting topic of discussion in future studies. Third, the regulation of immune processes, which influence growth by affecting general health. However, it should be kept in mind that the gene annotations by gene ontology have been based mainly based on the data on humans and the model animals (mouse, rat, etc.); thus, the true physiological roles of the identified genes in horses may vary. Also, the possible associations of variants that remained unannotated require future clarification with the updated annotation data for horse genomes.

## Conclusions

The Kazakh horses, with traditionally defined types and breeds, represent a single landrace with an absence of clearly expressed internal genetic structure. The traditional ways of horse breeding and husbandry in Kazakhstan have led to the formation of a relatively unspecialized landrace with genetic properties similar to a wild-living population. Along with the genetically similar Mongolian and Tuva breeds, the original genetic pool of Kazakh horses can potentially serve as a new source of genetic material for horse breeding, for the development of new breeds, or the improvement of existing lineages.

## Acknowledgments

The study was funded within the framework of the research project AP14870614 «Genetic marking of productive traits of the Kazakh horse of the Dzhabe type based on genome wide coverage SNP genotyping» (the Ministry of science and higher education of the Republic of Kazakhstan) and the targeted funding program BR10764999 “Development of technologies for effective management of the breeding process and preservation of the gene pool in horse breeding” (the Ministry of Agriculture of the Republic of Kazakhstan).

---

Supplementary table 1. Summary of horse sample collection.

Supplementary table 2. List of Kazakh horses identified as hybrids (outlying genotypes)

Supplementary table 3. Results of annotation of SNP associated with size and weight in the Kazakh horses.

Supplementary figure 1. Neighbor joining tree of the Kazakh horses in comparison with foreign breeds.

## Footnotes

1 Variants of transliteration: Jabe, Dzhabe.

2 Variants of transliteration: Aday, Adaev.

3 Variant of transliteration: Kustanai.

## Notes

### Competing Interest Statement

The authors have declared no competing interest.

### Summary of Updates

Correct the title

