## Supplementary figures and images for "Genetic structure and genome-wide association study of the traditional Kazakh horses"

### Supplementary figure 1

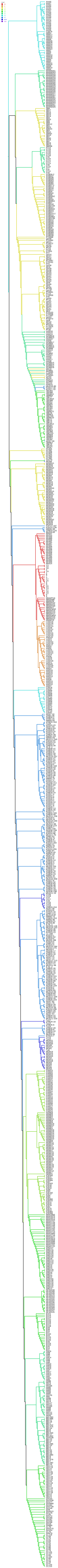
